# Blue light increases stomatal conductance and photosynthesis in *Agave* hybrid

**DOI:** 10.1101/2025.05.13.653724

**Authors:** Heitor Lopes Sartori, Sávio Justino da Silva, Larissa Prado da Cruz, Matheus Dalló Laira, Fábio Trigo Raya, Gonçalo Amarante Guimarães Pereira, Rafael Vasconcelos Ribeiro

## Abstract

Blue light (BL) plays an important role in stomatal opening, finely tuning plant responses to environmental conditions. While the BL signaling pathway is well understood in C_3_ and C_4_ plants, its role in crassulacean acid metabolism (CAM) plants remains uncertain. Traditionally, stomata in CAM plants were considered insensitive to BL stimulation, and as a result, studies on such interaction were overlooked for a long time. Only recently, studies have found that the BL signaling cascade is active in CAM plants. Here, we investigated the effects of BL intensity on stomatal behavior in *Agave*, a highly productive CAM plant, by stimulating *Agave* leaves with BL and taking measurements of leaf gas exchange during the morning (closed stomata) and in the afternoon (open stomata). Our findings revealed that BL had no significant effect on stomatal opening during the morning period. However, BL increased stomatal conductance (*g*_*s*_) by 67.3%, photosynthetic rate (*A*) by 109.5%, and intrinsic water use efficiency (iWUE) by 69.5% in the late afternoon. These findings suggest that *Agave* responded to BL similarly to C_3_ and C_4_ species when stomata were already open. We hypothesized that high CO_2_ levels due to C_4_ acid decarboxylation during the morning act as a repressive factor for stomatal opening by inhibiting the phosphorylative activity of the HT1 protein, a key protein involved in stomatal behavior. This inhibition likely prevents the activation of the signaling pathway mediated by CBC1/2 kinase, which integrates blue light (BL) signals with low intercellular CO_2_ levels.

## 1 Introduction

Stomata open and close in response to changes in external and internal plant signals (Auler et al., 2022) and such fine tuning of stomatal aperture ensures enough CO_2_ for photosynthesis while maintaining water balance and leaf temperature, which are critical for plant growth and survival (Brodribb et al., 2020; Santos et al., 2021). Light is one of the most important external factors stimulating stomatal opening, and this irradiance-mediated behavior occurs through two distinct signaling pathways: the “red” light (RL) – also known as photosynthetic/mesophyll – response; and the “blue” light (BL) response, which is independent on photosynthesis and specific to guard cells (GC) (Shimazaki et al., 2007). Despite these distinct pathways, both RL and BL can trigger phosphorylation of the plasma membrane H^+^-ATPase in guard cells, leading to membrane hyperpolarization, K^+^ uptake and subsequent stomatal opening (Ando and Kinoshita, 2018).

The mechanisms involved in RL pathway for inducing stomatal opening have been studied (Zhu et al., 2020). The relationship between RL and stomatal opening occurs indirectly, due to the reduction in internal CO_2_ concentration promoted by photosynthetic activity in mesophyll cells, as demonstrated by Roelfsema et al. (2002). Moreover, RL likely facilitates stomatal opening through an increase in turgor pressure in guard cells, which is driven by the accumulation of mesophyll-derived sucrose (Flütsch et al., 2020; Flütsch and Santelia, 2021). In parallel, BL also plays a critical role in stomatal regulation. BL activates the plasma membrane H^+^-ATPase pump through a signaling cascade triggered by the autophosphorylation of guard cell phototropins, which hyperpolarizes the plasma membrane, leading to the accumulation of osmolytes and promoting water influx into GCs, driving stomatal opening (Ando and Kinoshita, 2018; Bernardo et al., 2023). BL-dependent stomatal opening requires RL as background light source, with the most pronounced responses observed when both spectra are supplied simultaneously (Doi et al., 2015; Li et al., 2023).

However, the response pathway to BL has not yet been fully elucidated in crassulacean acid metabolism (CAM) plants. Although some studies claim that the BL-mediated signal transduction in GCs occurs predominantly in C_3_ and C_4_ plants and is not applicable to the stomatal movement of CAM plants (Tallman et al., 1997; Lee, 2010; Lee, 2019), Gotoh et al. (2019) demonstrated that the obligatory CAM species *Kalanchoe pinnata* and *K. daigremontiana* opened their stomata in response to BL. These results were obtained by exposing plants to light stimuli combining low BL and high RL photon flux densities (10 and 600 μmol m^−2^ s^−1^, respectively), with stomatal opening observed in both detached epidermis and intact leaves. Therefore, it becomes evident that further research is needed to clarify and establish the mechanisms involved in stomatal dynamics in CAM plants.

Unlike most CAM species, the *Agave* genus stands out as one of the few capable of achieving impressive productivities (Garcia-Moya et al., 2011). This makes it not only a valuable source for fibers and alcoholic beverages, but also a promising candidate for biofuel production in semiarid regions and as an alternative crop for areas most affected by the climate crisis (Raya et al., 2022). Additionally, due to its robustness and high tolerance to drought, there is growing interest in understanding CAM in agriculturally important species, with the aim of bioengineering this pathway into C_3_ plants to enhance water use efficiency in traditional crops (Borland et al., 2014; Yang et al., 2015; Cushman et al., 2019; Lim et al., 2019). In *Agave*, blue-light-responsive genes exhibit a temporal expression pattern similar to that of a C_3_ plants (Abraham et al., 2016). However, the responsiveness of these plants to BL has never been tested. Also, BL supplementation would be a strategy for enhancing photosynthesis and growth (Zheng et al., 2018; Kaiser et al., 2019; Moradi et al., 2021). *Agave* cultivation could also benefit from such light management, especially during seedling production stages, potentially accelerating growth in greenhouses and increasing the supply of seedlings needed to achieve large-scale biofuel production.

Here, the effects of BL on the modulation of stomatal behavior in *Agave* were explored, providing valuable insights into how these species respond to light stimuli and opening new possibilities for optimizing their early cultivation in controlled environments.

## 2 Material and methods

### 2.1 Plant material and growing conditions

Bulbils from the *Agave* hybrid “Four-Hundred Leaves”, a prominent Brazilian cultivar of unknown origin (De Souza et al., 2018; Raya et al., 2023), deposited in Unicamp’s germplasm bank under accession LBF1 were cultivated for nine months in a greenhouse in Campinas, São Paulo, Brazil (22°49’S, 47°4’W), under average air temperature and relative humidity of 24.2 ± 2°C and 73 ± 9%, respectively (Supplementary Material Fig. S1A). The plants were grown in 1.5 L plastic pots filled with a mixture of sand, organic compost, and vermiculite (3:2:1 ratio). They were fertilized every ten days with full-strength Sarruge nutrient solution (Sarruge, 1975) and irrigated every three days.

### 2.2 Blue light variation and leaf gas exchange

Measurements were conducted on the fourth leaf (counting from the central spike) of five individual plants. For each plant, data were collected at two time points: once in the morning, when stomata were closed (10h00–12h00) and once in the afternoon, when they are open (16h00–18h00). This resulted in two measurements per plant per treatment. We used a porometer (model LI-600PF, LI-COR, Lincoln NE, USA) to follow the daily pattern of stomatal opening inside the greenhouse, ensuring accurate timing. Due to the number of treatments, multiple days were required to complete all measurements under controlled conditions.

Plants were moved to laboratory and maintained under room conditions (average air temperature and relative humidity of 25.4 ±1°C and 71 ± 3%, respectively). The proportion of BL (λ = 460 nm) and RL (λ = 635 nm) incident on leaves were controlled and leaf gas exchange measured using an infrared gas analyzer (LI-6400XT, LI-COR, Lincoln NE, USA), equipped with a modulated fluorometer (model 6400-40, LI-COR, Lincoln NE, USA) (Supplementary Material Fig. S1B,C). Measurements were taken under air CO_2_ concentration of 400 µmol mol^-1^ and incident photosynthetically active radiation (PAR) of 430 μmol m^−2^ s^−1^. Leaves were clamped to the device until stabilization of leaf gas exchange before beginning the measurements to ensure any variation was due to varying BL proportion. Blue/red light ratios of 10/90, 50/50, 90/10, 0/100, and 10/90 were applied sequentially, corresponding to BL/RL photon flux densities of 43/397, 215/215, 397/43, 0/430, and 43/397 μmol m^−2^ s^−1^, respectively. Each light treatment was applied for 30 minutes, during which the net photosynthetic rate (*A*), stomatal conductance (*g*_*s*_), *A*/*g*_*s*_ratio – intrinsic water use efficiency – (iWUE), intracellular CO_2_ concentration (C_i_) and transpiration (*E*) were recorded every minute along each blue light step. Since all light regimes were applied for the same duration and under constant photon flux density, the total fluence (i.e., cumulative photon dose) varied proportionally with each light treatment.

To account for the natural variability in absolute values among biological replicates and to better visualize treatment-induced trends, the data for *g*_*s*_, *A*, iWUE, C_i_ and *E* were normalized (*n’*, where *n* is the parameter analyzed) according to Equation (1), where *n*_*min*_and *n*_*max*_represent the minimum and maximum values obtained for each parameter.

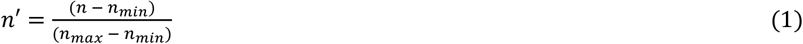

The area under the curve (AUC) for *g*_*s*_, *A*, and iWUE was calculated to determine the gains when varying blue light proportions. Calculations were based on the trapezoidal integration method using the “*trapz”* function of the “*pracma”* package in R (R Core Team, 2024). The method approximates the integral of the curve as the sum of trapezoidal areas between adjacent points, as described in Equation (2):

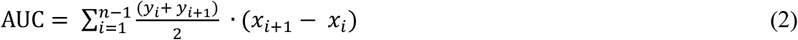

where, *x*_*i*_ and *x*_*i*+1_ are consecutive time points, and *y*_*i*_ and *y*_*i*+1_ are the corresponding parameter values. The difference between AUC of plants exposed to varying BL proportions (AUC_*BLv*_, Fig. 1) and the AUC of plants maintained under constant BL (AUC_*BLc*_, Supplementary Material Fig. S2) represents the gain achieved by supplying BL for each photosynthetic parameter, as detailed in Equation (3):

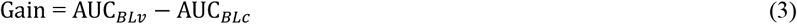

**Fig. 1.**
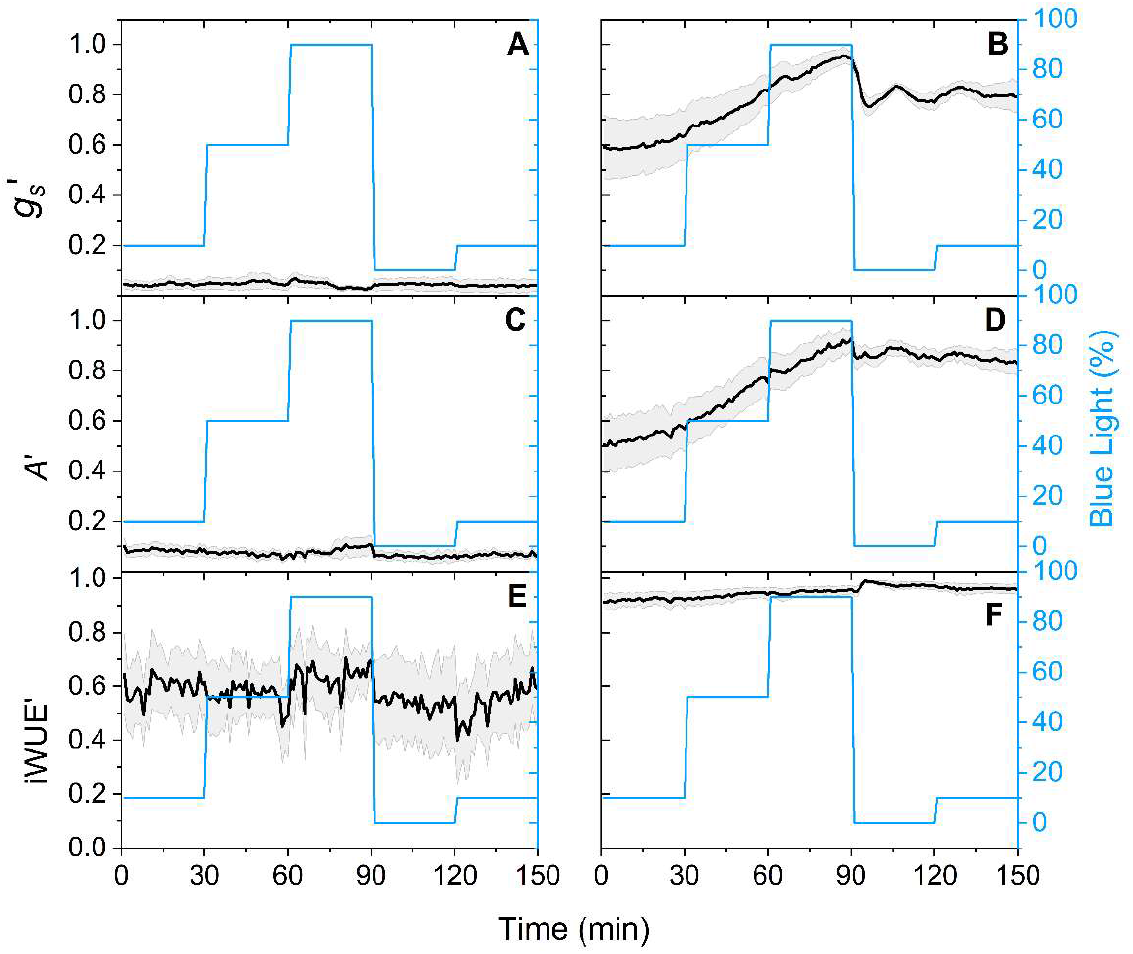
Time-course of normalized stomatal conductance (*g*_*s*_’), net photosynthetic rate (*A*’), and intrinsic water use efficiency (iWUE*’*) in *Agave* plants under varying blue/red light ratios of 10/90, 50/50, 90/10, 0/100, and 10/90 (corresponding to blue/red photon flux densities of 43/397, 215/215, 397/43, 0/430, and 43/397 μmol m^−2^ s^−1^, respectively). Measurements were taken during the morning (A, C, and E) and afternoon (B, D, and F). Black lines represent the mean values (n = 5) and the shaded areas are the standard error.

### 2.3 Data analysis

The results were analyzed using the F-test, and regression analysis was done when appropriate (software Origin 9.1, OriginLab Corporation, Northampton MA, USA). Mean values from boxplots were also analyzed using Bayesian statistics, with comparisons made through the Bayes Factor (BF_10_): BF_10_ < 1 favors the null hypothesis (*H*0); 1 ≤ BF_10_ < 3 suggests weak evidence for the alternative hypothesis (*H*1); 3 ≤ BF_10_ < 20 indicates moderate evidence for *H*1; 20 ≤ BF_10_ < 150 suggests strong evidence for *H*1 and BF_10_ ≥ 150 indicates very strong evidence for *H*1 (Kass and Raftery, 1995).

## 3 Results and discussion

In the morning, when the stomata were closed, varying proportions of BL did not result in any noticeable change in leaf gas exchange (Fig. 1A,C,E and Supplementary Material Fig. S3A,C). The normalized mean values during the evaluation period remained steady at approximately 0.046 for *g*_*s*_′ and 0.079 for *A*′, corresponding to absolute values of 0.0035 mol m^−2^ s^−1^ and -0.45 μmol m^−2^ s^−1^, respectively. These findings are consistent with expectations and align with previous studies on CAM plants, which have reported similar responses when exposed to BL (Lee and Assmann, 1992; Tallman et al., 1997; Kinoshita; Hayashi, 2011). For a long time, these findings reinforced the assumption that CAM plants were “unresponsive” to BL, suggesting that stomatal opening mechanisms were not directly influenced by it. However, during the late afternoon and when stomata were already open, varying BL proportions caused significant changes in leaf gas exchange (Fig. 1B,D,F and Supplementary Material Fig. S3B,D). Specifically, we observed increases of approximately 67.3% (from 0.049 to 0.082 mol m^−2^ s^−1^) in *g*_*s*_, 109.5% (from 4.3 to 9.0 μmol m^−2^ s^−1^) in *A*, and 69.5% in iWUE. Compared to plants maintained under constant BL (Fig. S2) and considering AUC values, varying BL proportions caused significant gains in all analyzed photosynthetic parameters: +92.3% in *g*_*s*_, +51.6% in *A*, and +92.6% in iWUE. The increase in intrinsic water use efficiency (iWUE) suggests an enhanced capacity of the plant to fix carbon per unit of water lost. This response is particularly relevant for species adapted to arid environments, such as *Agave*, as it may reflect a blue-light-mediated optimization of the balance between CO_2_ uptake and water conservation. Circadian rhythms may also influence stomatal responsiveness to blue light in *Agave*, as varying behaviors were observed between morning and afternoon. Although not the focus of this study, future investigations on how circadian regulation interacts with blue light signaling could offer deeper insights into stomatal control in CAM plants.

The BL-induced stomatal opening was further confirmed by the simultaneous decrease in *g*_*s*_and *A* after 90 minutes, when the BL proportion dropped from 90% to 0% (Fig. 1B,D). This shift led to 18.3% reduction in *g*_*s*_, changing from 0.082 to 0.067 mol m^−2^ s^−1^. Contrary to prior assumptions, these results confirm the presence of BL-responsive mechanisms in CAM plants, as first indicated by Ceusters et al. (2014) in *Aechmea ‘Maya’*, where low-fluence BL played a key role in stomatal regulation. Our findings also align with those of Gotoh et al. (2019), who demonstrated that H^+^-ATPase is involved in the stomatal opening signal transduction pathway in CAM plants in response to BL, similar to C_3_ and C_4_ plants. In a broad perspective, our findings and the current literature suggests that plant mechanisms governing BL response are independent on the photosynthetic model. Furthermore, plants were unable to restore the initial *g*_s_ and *A* values when providing 100% red light (Fig. 1B,D), implying that guard cells may retain a memory to BL, taking longer to fully reverse its effects (Lüttge and Thellier, 2016).

The significantly positive linear regression (p<0.05) between leaf gas exchange parameters and BL proportion when the plants had open stomata indicates that stomatal conductance, photosynthesis and intrinsic water use efficiency are modulated by BL, with the lowest values observed in plants exposed to 10% BL and the highest values in plants exposed to 90% BL during the experimental period (Fig. 2). Here, the time step of 30 min was not enough to stabilize leaf gas exchange, e.g., stomata was progressively opening after changing BL proportion (Fig. 1B,D). In fact, the extent of the transient stomatal opening induced by BL may vary depending on the species and experimental conditions (Zhen and Bugbee, 2020), such as the level of background RL (Suetsugu et al., 2014). Here, the increase in leaf gas exchange when increasing the proportion of BL suggests that the signaling pathway specific to GCs in CAM plants has a greater influence on stomatal behavior than the mesophyll signaling pathway. Anyway, our data are consistent with the hypothesis that BL activates a metabolic signaling cascade that regulate stomatal opening in *Agave*, which opens new possibilities for BL application in controlled production systems, such as biofactories or nurseries. As stomatal opening and photosynthesis are closely linked to plant growth rate, one would enhance biomass production by providing BL to plants.

**Fig. 2.**
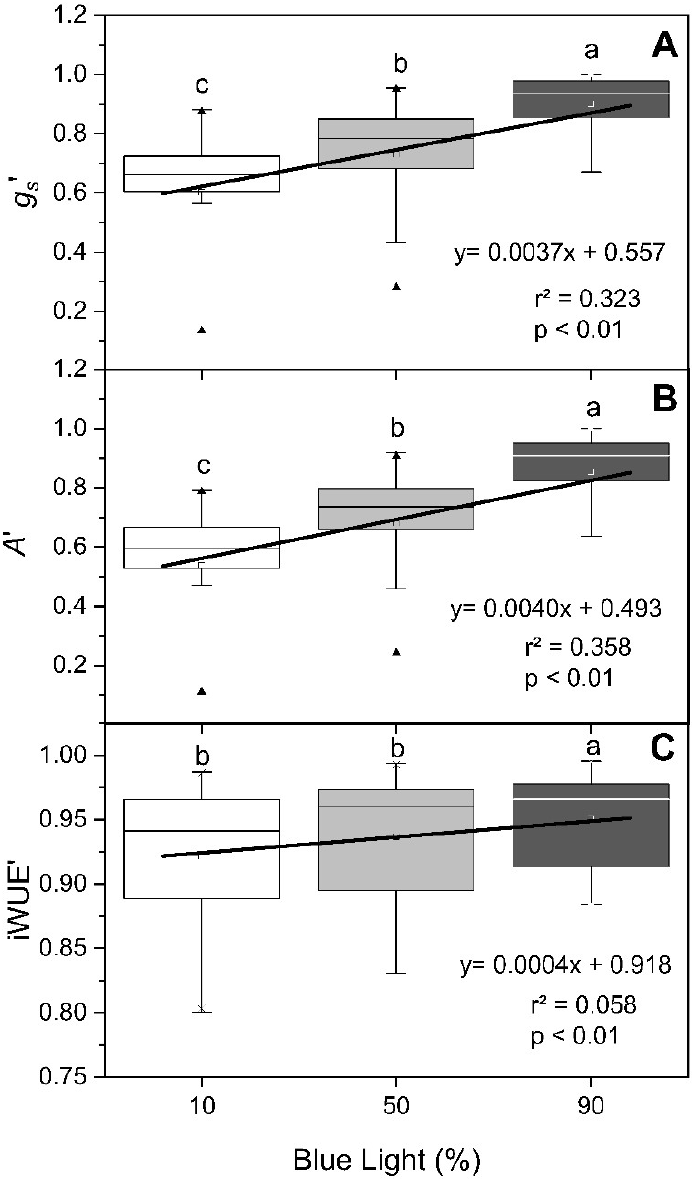
Normalized stomatal conductance (*g*_*s*_’, in A), net photosynthetic rate (*A*’, in B), and intrinsic water use efficiency (iWUE*’*, in C) in *Agave* plants as affected by blue light proportion supplied at afternoon. Different letters above the boxplots indicate statistically significant differences between BL proportions (BF_10_ > 150). Each boxplot is composed by measurements taken during 30 min (n = 150). Blue light proportion is complemented with red light, e.g., 10% blue light (λ = 460 nm) and 90% red light (λ = 635 nm). The black line in each panel represents the linear regression model with corresponding equation, r^2^ value and *p*-value.

Additionally, our results suggest a repressive factor in the morning that inhibits stomatal opening despite BL stimulation. As the intercellular CO_2_ concentration (*C*_i_) decreased progressively in the afternoon when blue light proportion increased, we have evidence that *C*_i_ dynamics might play a role in modulating stomatal aperture (Fig. 2B and Fig. S3B). One possible hypothesis is that this repression is linked to the intercellular CO_2_ concentration. During the CAM cycle, malic acid is decarboxylated throughout the day, generating the CO_2_ necessary for photosynthesis. It is reasonable to propose that once the CO_2_ reserves are depleted, the stomata open. Recently, Hiyama et al. (2017) identified and characterized a novel protein kinase called CBC1/2 (CONVERGENCE OF BLUE LIGHT (BL) AND CO_2_ 1/2), which stimulates stomatal opening by integrating BL signals with low CO_2_ levels. The activity of this protein depends on phosphorylation induced by HT1, a Raf-like protein kinase that plays a critical role in the phosphorylation of proteins involved in stomatal behavior (Hashimoto et al., 2006; Takahashi et al., 2022). Early in the day when CO_2_ availability in intercellular spaces is high due to decarboxylation of malate, CBC1/2 remains dephosphorylated and, consequently, inactive, because high concentrations of CO_2_ inhibit HT1 activity (Takahashi et al., 2022). Most studies evaluating the signaling pathway mediated by CBC1/2 have been conducted in *Arabidopsis*, but our findings suggest that this mechanism must be robust and conserved in the *Agave* genus for CAM metabolism to function efficiently.

Lastly, blue light has been associated with changes in starch and lipid metabolism in *Arabidopsis* guard cells, contributing to stomatal opening. Starch degradation provides carbon skeletons for the synthesis of malate, whose accumulation leads to increased turgor and subsequent stomatal opening (Flütsch et al., 2020). Additionally, triacylglycerol mobilization supplies the ATP required for this process (McLachlan et al., 2016). Although these mechanisms are well described in C3 species, there is currently no clear evidence of their role in CAM plants such as *Agave* spp. To address these questions, further research is crucial to deepen our understanding on the mechanisms that modulate stomatal opening in response to BL, especially in CAM plants, where the interplay between light and other factors remains underexplored.

## Supporting information

BlueLight_SupplementaryMaterial

## Acknowledgments

This work was supported by the Agência Nacional do Petróleo, Gás Natural e Biocombustíveis do Brasil (ANP) in association with Shell Brasil Petróleo Ltda, with investments from the PD&I Clauses, for the Brazilian *Agave* Development Program (BRAVE) [Research agreements 5902 and 5903]. The authors are grateful to LCroP and LGE members for the insightful discussion during our group meetings. Rafael V. Ribeiro is a fellow of the National Council for Scientific and Technological Development [CNPq, Brazil, grant no. #304295/2022-1].

## Data availability

Data will be made available if requested to the corresponding author.

