## Supplementary material for "Blue light increases stomatal conductance and photosynthesis in *Agave* hybrid": BlueLight_SupplementaryMaterial

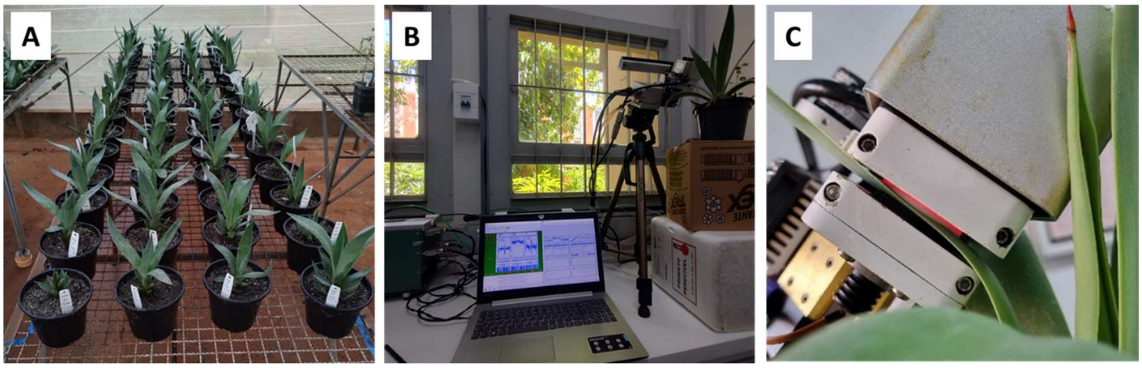


**Fig. S1** Experimental setup. Nine-month-old *Agave* plants cultivated in the greenhouse (A), measurement of a plant using the LI-6400XT system in the laboratory (B), and positioning of the *Agave* leaf inside the LI-6400XT chamber (C).

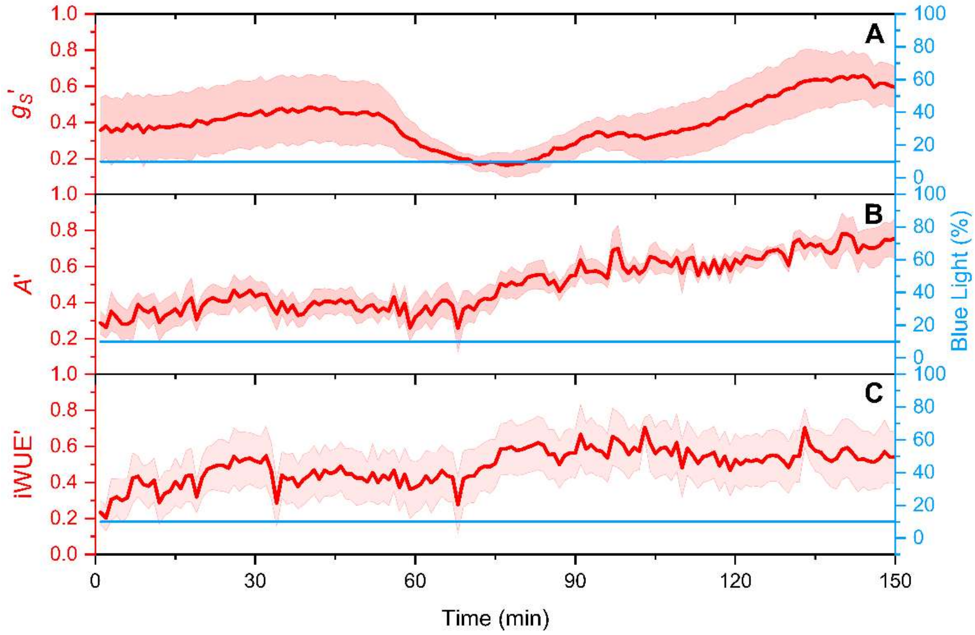


**Fig. S2** Time-course of normalized stomatal conductance ($g_s$'), net photosynthetic rate ($A$'), and intrinsic water use efficiency (iWUE') in *Agave* plants exposed to constant blue/red light ratio of 10/90 (corresponding to blue/red intensities of 43/397 μmol m$^{-2}$ s$^{-1}$, respectively). Measurements were taken during the afternoon. Red lines represent the mean values (n = 5) and the shaded areas are the standard error.

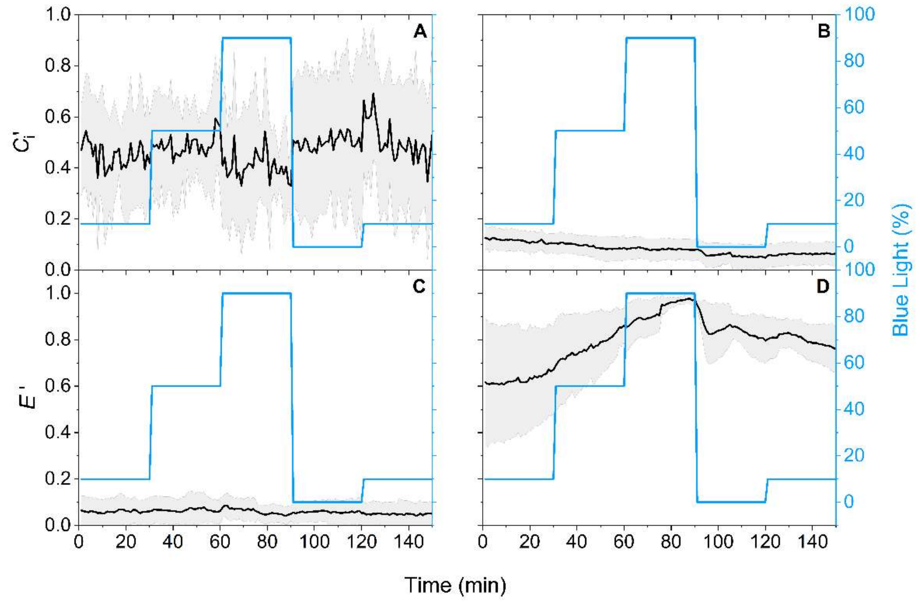


**Fig. S3** Time-course of normalized intracellular $CO_2$ concentration ($C_i$') and transpiration (*E*') in *Agave* plants under varying blue/red light ratios of 10/90, 50/50, 90/10, 0/100, and 10/90 (corresponding to blue/red photon flux densities of 43/397, 215/215, 397/43, 0/430, and 43/397 µmol $m^{-2}$ $s^{-1}$, respectively). Measurements were taken during the morning (A and C) and afternoon (B and D). Black lines represent the mean values (n = 5) and the shaded areas are the standard error.
